# δ-α cell-to-cell interactions modulate pancreatic islet Ca^2+^ oscillation modes

**DOI:** 10.1101/2024.08.21.608986

**Authors:** Huixia Ren, Yanjun Li, Beichen Xie, Weiran Qian, Yi Yu, Tianyi Chang, Xiaojing Yang, Kim Sneppen, Liangyi Chen, Chao Tang

## Abstract

Glucose-induced pancreatic islet hormone release is tightly coupled with oscillations in cytoplasmic free Ca^2+^ concentration of islet cells, which is regulated by a complex interplay between intercellular and intracellular signaling. δ cells, which entangle with α cells located at the islet periphery, are known to be important paracrine regulators. However, the role of δ cells in regulating Ca^2+^ oscillation pattern remains unclear. Here we show that δ-α cell-to-cell interactions are the source of variability in glucose-induced Ca^2+^ oscillation pattern. Somatostatin secreted from δ cells prolonged the islet’s oscillation period in an α cell mass-dependent manner. Pharmacological and optogenetic perturbations of δ-α interactions led islets to switch between fast and slow Ca^2+^ oscillations. Continuous adjustment of δ-α coupling strength caused the fast oscillating islets to transition to mixed and slow oscillations. We developed a mathematical model, demonstrating that the fast-mixed-slow oscillation transition is a Hopf bifurcation. Our findings provide a comprehensive understanding of how δ cells modulate islet Ca^2+^ dynamics and reveal the intrinsic heterogeneity of islets due to the structural composition of different cell types.

**Highlights:**

- Somatostatin slows down islet Ca^2+^ oscillations in an α cell mass-dependent manner.
- Pharmacological and optogenetic perturbations of δ-α interaction cause islet Ca^2+^ oscillation mode switching.
- Continuous tuning of δ-α interaction strength induces fast-mixed-slow oscillation transition successively.
- Mathematical modeling shows the fast-mixed-slow oscillation transition as a Hopf bifurcation.

## Introduction

An organ consists of multiple cells of different types, the interaction among which is critical for the complex function it performs. Understanding how the different cell types coordinate to orchestrate the organ function is also crucial for revealing the cause of dysfunction in these multicellular networks^1^. Pancreatic islets are a prototypical example of such a system, whose function is to regulate the blood sugar level via the secretion of hormones (insulin and glucagon)^2^.

A pancreatic islet is mainly comprised of three cell types: β, α and δ, which make up approximately 80%, 15% and 5% of the total cell population, respectively^3^. Functionally, hormone secretion from an islet cell is closely coupled to the oscillation in the cell’s cytoplasmic free Ca^2+^ concentration^4–6^. Various Ca^2+^ oscillation patterns have been observed in islets, including fast (∼20 seconds) oscillation, slow (∼5 minutes) oscillation and mixed oscillations of fast and slow^7–9^. Frequency-modulated oscillations of cytosolic Ca^2+^ are thought to play a crucial role in insulin secretion^6^, and their dysfunction is associated with aging^10^ and diabetes^11–13^.

In an islet, the insulin secreting β cells are extensively coupled via connexin36 (Cx36) gap junction and display a more homogeneous response^14^. Notably, regular fast oscillation of β cells mainly exists in intact islets^15^. In isolated single β cells, the Ca^2+^ oscillation is typically slow with irregular fast and display high heterogeneity^9,15,16^. The differences in behavior between cells in isolated and intact islet environment highlight the importance of cell-cell interactions within the islet.

The α cells secrete glucagon which can stimulate islet cells’ hormone secretion by elevating their cAMP level through G_s/q_ signaling^17^. Pharmacological studies found that glucagon accelerates the oscillation frequency in both islet^4,18,19^ and isolated β cells^20^ by increasing β cell’s cytosolic cAMP level and facilitating its endoplasmic reticulum Ca^2+^ release. Supporting this finding, islets in SERCA3 (ATPase sarcoplasmic/endoplasmic reticulum Ca^2+^ transporting 3) knockout mice lost fast Ca^2+^ oscillation^21^. The δ cells exhibit dendrite-like structure which can communicate with many neighboring α and β cells to act as efficient regulators of activity^22^. δ cells secrete somatostatin, which inhibits insulin secretion from β cells by decreasing their cAMP level through G_i_ signaling^23^. Somatostatin also inhibits glucagon secretion from α cells by activating G_i_-coupled Sstr2^24–26^. Somatostatin knockout mice exhibit significantly elevated basal and glucose-stimulated glucagon secretion^27^. Moreover, the short bioactive half-life of somatostatin suggests its inhibitory action through paracrine signaling^22^. These findings suggest that islet’s oscillation frequency may depend on the paracrine interactions in islets.

In this study, we investigate the critical role of δ-α cell-to-cell interaction in modulating the islet Ca^2+^ oscillation modes. We used a microfluidic platform which provides a controllable and stable perfusion environment for high-throughput imaging of intact islets. In combination with the α cell-labeled transgenic mice, this platform allowed for a systematic investigation for our purpose. We found that α cells are key to inducing fast Ca^2+^ oscillations, while δ cells are crucial for recovering β cells’ slow Ca^2+^ oscillations by inhibiting α cells. Pharmacological and optogenetic perturbations confirmed that δ-α cell interactions generate tunable islet oscillation patterns. Furthermore, continuous parameter tuning demonstrated that balanced regulations of α and δ cells could induce mixed (fast and slow) Ca^2+^ oscillations. Hopf bifurcation refers to the appearance or disappearance of a periodic solution from an equilibrium as a control parameter crosses a critical value. Combined with a mathematical model, we showed that the transition is a Hopf bifurcation resulting from the paracrine interaction between α and δ cells.

## RESULTS

### High-throughput Imaging of Ca^2+^ Activity and 3D Structural Composition of Islets

Simultaneous imaging of islet Ca^2+^ activity and its 3D cell composition enables investigation of the islets’ intrinsic heterogeneity due to structural composition of different cell types. To achieve this goal in a high-throughput manner, we designed a microfluidic chip (Fig. 1a and Methods). This device enabled the simultaneous comparison of eight islets from two sets of observations, each consisting of a flexible combination of up to five perfusion solutions. To determine the cell composition in islets, we generated a triple transgenic mouse line (*Glu-Cre^+^; tdTomato^f/+^; GCaMP6f^+^*) by crossbreeding of *GCaMP6f^+^ and Glu-Cre^+^; tdTomato^f/f^* mice. Using two-photon microscopy, we identified individual α cells and measured the islet diameter while recording the islet’s Ca^2+^ signal using spinning disk microscopy (Fig. 1b-c and Methods).

**Fig. 1.**
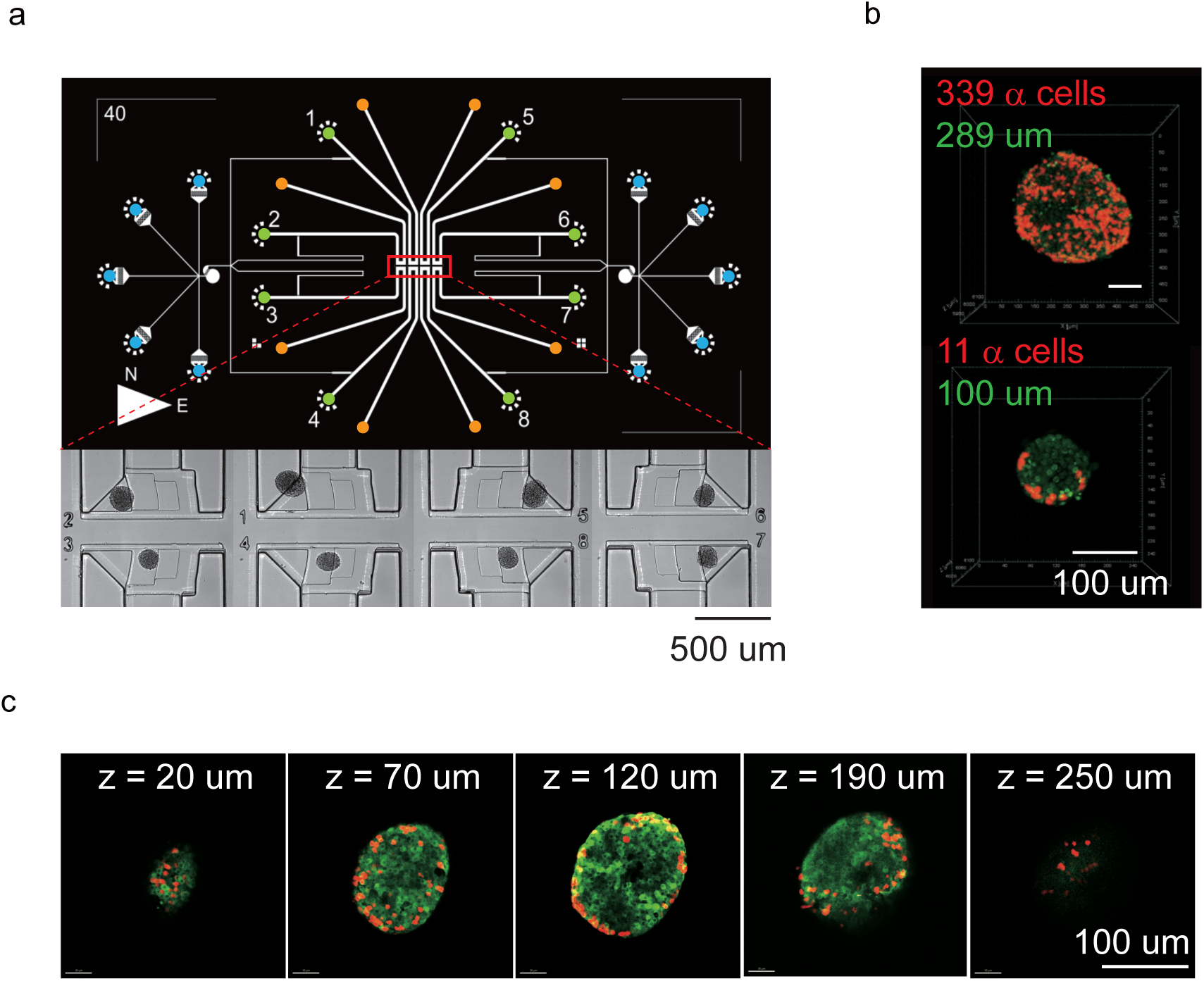
Microfluidic chip for imaging multiple islets. a) Upper panel: Illustration of the microfluidic chip designed for islet Ca^2+^ imaging. The red box indicates the imaging zone. The blue circular ports serve as reagent inlets, the green circular ports function as islet inlets, and the yellow circular ports serve as reagent outlets. Bottom panel: A bright-field image displaying the imaging zone, where 8 islets have been loaded. b) 3D projection of *Glu-Cre^+^; tdTomato^f/+^; GCaMP6f^+^* islets. α cells expressing tdTomato proteins are highlighted in red, while all islet cells expressing GCaMP6f protein are shown in green. The red number indicates the count of α cells, and the green number represents the islet diameter. c) Z-stack images of *Glu-Cre^+^; tdTomato^f/+^; GCaMP6f^+^* islets. The number indicates the depth.

### Glucagon Accelerates and Somatostatin Slows Down Ca^2+^ Oscillation

Under high glucose stimulation, pancreatic islets exhibit a conserved and α cell mass dependent activity pattern. The first response to the elevated glucose concentration from 3 to 10 mM (3G to 10G) was a pronounced Ca^2+^ peak followed by regular oscillations, which were slow in islets with <40 α cells (Figs. 2a-c, 100-400 s per cycle, see Methods for detailed oscillation spectrum analysis) and fast in the remaining islets (10-40 s per cycle). Furthermore, adding glucagon (100 nM) to the media uniformly transformed islets’ activity into fast Ca^2+^ oscillations, regardless of the number of α cells (Fig. 2c). In contrast, somatostatin (1 µM) slowed down the Ca^2+^ oscillations in islets with <100 α cells. However, α-cell abundant islets maintained fast Ca^2+^ oscillations at 1 µM somatostatin concentration. Moreover, the order of adding glucagon and somatostatin did not affect the results (Fig. 2b). It is worth noting that the oscillation periods of the islets before and after the perturbation followed a bimodal distribution. While without/with perturbation larger islets tended to oscillate faster, there were no significant differences (Fig.2d). These findings demonstrated that glucagon and somatostatin could have a dominating influence on the glucose-stimulated islet’s Ca^2+^ oscillation mode.

**Fig. 2.**
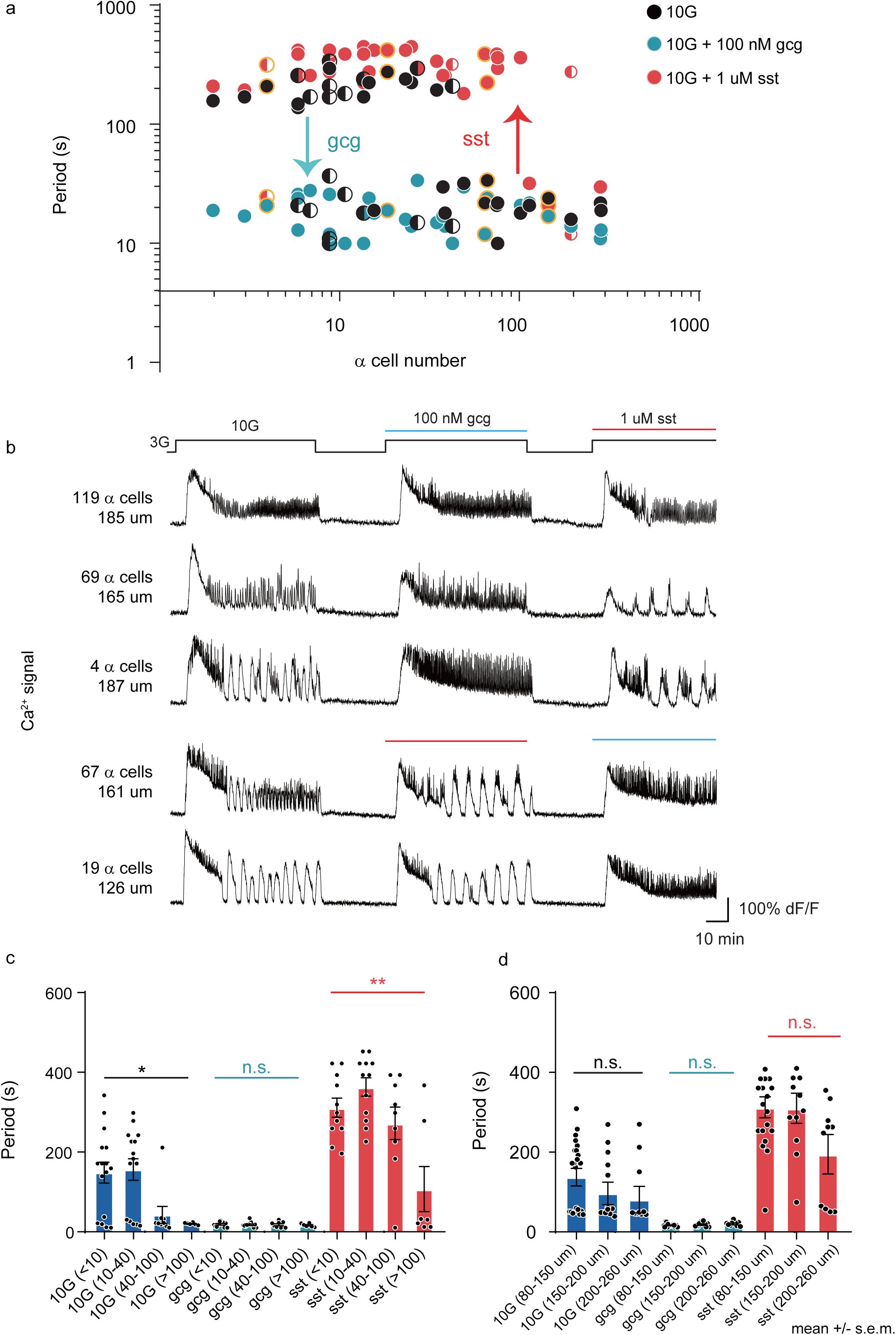
Glucagon and Somatostatin Switch Ca^2+^ Oscillation mode between Slow and Fast. a) Scatter plot of islet’s α cell number (tdTomato positive cell number) and mean islet Ca^2+^ oscillation period under 10G stimulation (black dots), 100 nM glucagon (blue dots) and 1 uM somatostatin (red dots) (n = 37 islets from 5 mice in 5 independent isolations). Half-filled dots indicate the islets with mixed oscillations. The islets labeled with yellow edge dots were shown in (b). b) Mean islet Ca^2+^ traces with 3G and 10G stimulation, and in addition of 100 nM glucagon (blue) and 1 uM somatostatin (red). The number of α cells (tdTomato positive cells) and the diameter of the islet are shown on the left. c) Ca^2+^ oscillation periods for islets with <10, 10-40, 40-100 and >100 α cells. Quantification of data in a). Bars represent mean ± s.e.m. (standard error of mean). d) Ca^2+^ oscillation period for islets diameter 80-150 um, 150-200 um and 200-260 um. Asterisks denote significance as 0.12 (n.s.), 0.0332 (*), 0.0021*(**), 0.0002(***) and <0.0001(****).

### Glucagon-Elevated cAMP Blocks Somatostatin-Induced Slow Ca^2+^ Oscillation

Somatostatin tuned islet Ca^2+^ oscillation in an α cell mass dependent manner, which suggests a kind of integration of glucagon and somatostatin pathways in the β cells. We further asked whether glucagon can block somatostatin’s modulation of β cells. Indeed, islets undergoing somatostatin-mediated slow Ca^2+^ oscillation showed fast Ca^2+^ oscillation when treated with 100 nM glucagon (Figs. 3a-b). Glucagon activates Gs/q-coupled GLP1Rs (Glucagon-like peptide-1 receptors) and GCGRs (glucagon receptors) to maintain β cell’s cAMP level^17^. Thus, we investigated whether glucagon blocks somatostatin’s modulation of β cell by elevating its cAMP level. Glucagon-treated islets exhibited elevated whole islet cAMP levels, while somatostatin-treated islets displayed decreased cAMP levels (Fig. 3c). Treatment with 12 µM forskolin resulted in fast Ca^2+^ oscillation in islets undergoing somatostatin-mediated slow Ca^2+^ oscillation (Figs. S2e-f, 287 s for 10G with somatostatin and 18 s for 10G with somatostatin and forskolin). These results suggest that glucagon and somatostatin pathways converge to control β cell cAMP level in opposing ways to tune the Ca^2+^ dynamics.

**Fig. 3.**
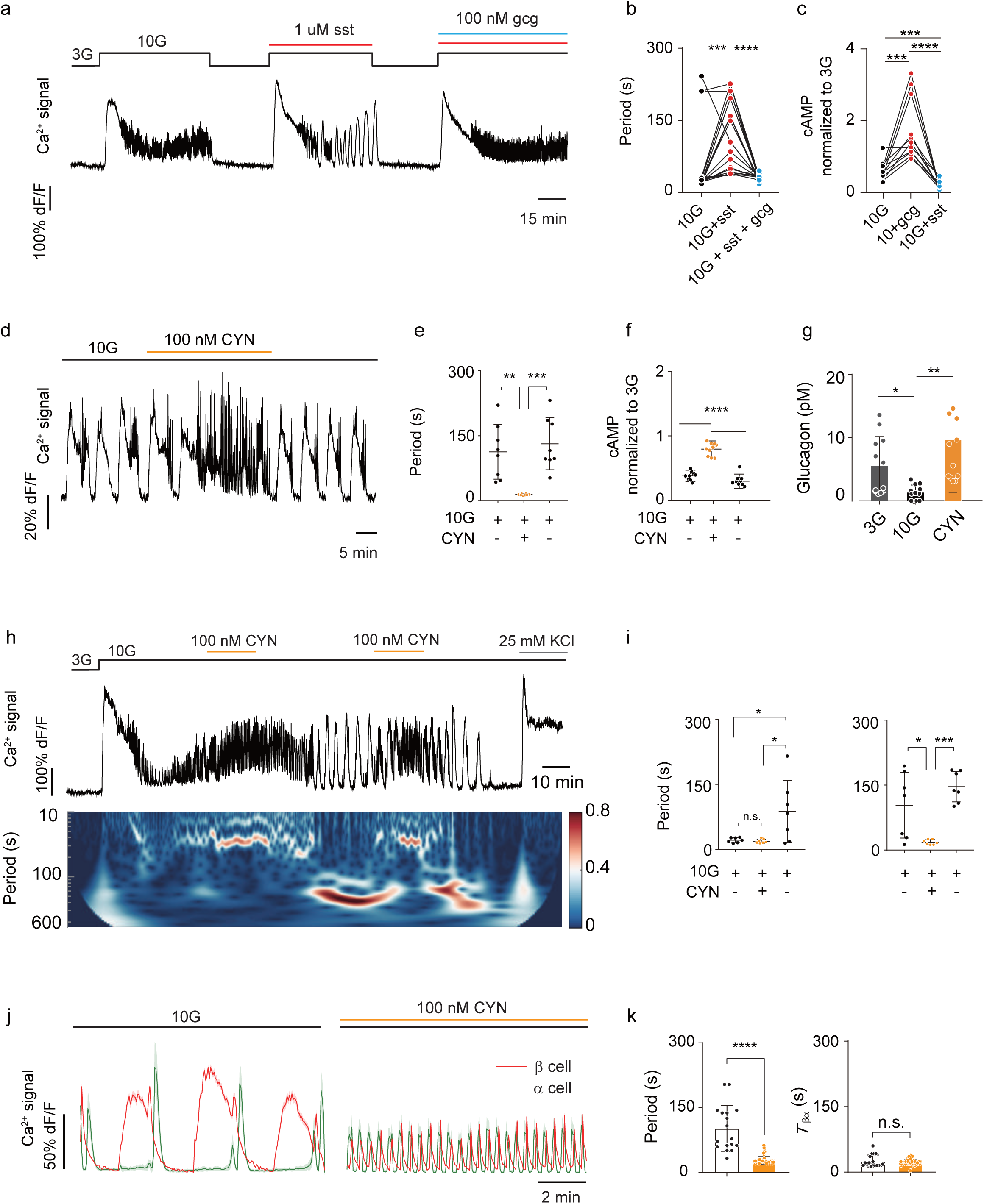
Somatostatin Limits Endogenous Glucagon Secretion and Slows Down Islet Ca^2+^ Oscillation. a) Recordings of whole islet Ca^2+^ signal with 10G, 1 uM somatostatin and 100 nM glucagon stimulation. Representative mean Ca^2+^ trace of whole islet under the stimulations of 10G, 1 uM somatostatin, 100 nM glucagon in present at 1 uM somatostatin, sequentially. b) Oscillation period before treatment (black, mean period 58 s), after somatostatin (red, mean period 150 s), and after somatostatin + glucagon (blue, mean period 31 s). Data represents n = 18 islets from 4 mice). c) Mean cAMP levels under different conditions: with 10G (black, 60%, normalized against the mean 3G level), 100 nM glucagon (blue, 174%), and 1 uM somatostatin (red, 20%). Data represents n = 11 islets from 3 C57BL/6J mice. d) Recordings of whole islet Ca^2+^ signal with 10G and 100 nM CYN stimulation (slow islets). e) Oscillation period before treatment, after CYN and washing out (mean 10G period 113 s, mean CYN period 14 s, and washout mean 10G period 132 s, n = 8 islets from 4 mice). Error bars in all panels depict SDs. f) Mean cAMP levels observed with different conditions: 10G (black, 38%, normalized against the mean 3G level), 100 nM CYN (yellow, 80%), and washing out (red, 30%). Data is based on n = 10 islets from 4 C57BL/6J mice. g) Secreted glucagon concentration for islets under 3G (grey, 5.6 pM), 10G (black, 1.3 pM) and 100 nM CYN (yellow, 9.6 pM). Data is based on n = 4 islets from 3 C57BL/6J mice (each islet measured for 30 min, 10 min per collection). h) Top: Recordings of whole islet Ca^2+^ signal (fast islets). The perfusion protocol was 10 min 3G, 40 min 10G, 20 min 100 nM CYN, 40 min 10G, 20 min 100 nM CYN, 40 min 10G and 15 min KCl. Bottom: Continuous wavelet transformation (CWT, see Methods) of whole islet Ca^2+^ signal. The Y-axis shows the CWT coefficients for periods ranging from 10 to 600 s. The color bar codes the CWT coefficients. i) Scatter plots of islet Ca^2+^ oscillation period before and after washout of CYN. Left: 1^st^ round of CYN added before mode switch to slow oscillation (10G period 21 s, CYN period 19 s, n = 7 islets from 3 mice). Right: 2^nd^ round of CYN added after mode switch (10G period 104 s, 100 nM CYN period 18 s, and washout 10G period 147 s, n = 7 islets from 3 mice). j) Recordings of mean Ca^2+^ activity of α (green) and β (red) cells under 10G stimulation without and with CYN (10 min) in microfluidic chip. The islets were isolated from *Glu-Cre^+^; GCaMP6f^f/+^*; *Ins2*-*RCaMP1.07* mice. Shadow indicates the standard deviation of each cell type. k) Period, waiting time of α cells (*T*_βα_) from experiments under 10G stimulation with and without CYN treatment (10G period was 104 s, 100 nM CYN period was 28 s, n = 17 (10G) and 76 cycles (CYN) from 4 islets in 3 *Glu-Cre^+^; GCaMP6f^f/+^*; *Ins2*-*RCaMP1.07* mice).

### δ Cell Limits Glucagon Secretion from α Cell and Slows Down Ca^2+^ Oscillation

δ cell is essential for slow Ca^2+^ oscillation and α cell for fast. We next set out to determine whether δ cells reduce the islet Ca^2+^ oscillation frequency via inhibiting α cells. Sstr2 is exclusively expressed in α cells and is required for somatostatin mediated glucagon secretion inhibition^24–26^. Therefore, we assessed how selective blockage of Sstr2 influenced islets showing 10G-stimulated slow and fast Ca^2+^ oscillations. In the presence of selective Sstr2 antagonist CYN-154806^28^ (CYN, 100 nM), islets showing slow Ca^2+^ oscillation transformed to fast oscillation, and recovered to slow oscillation after washout (Figs. 3d-e). For islets showing fast oscillation, CYN perfusion did not significantly change the period (Figs. 3h and 3i). Interestingly, after washout, these islets spontaneously switched into slow oscillation in approximately 30 minutes (Figs. 3h and 3i, washout 10G period 87 s). These data indicated that the glucose-stimulated Ca^2+^ oscillation mode may spontaneously change in a same islet. Either depletion or insufficient glucagon secretion can cause spontaneously slowdown. We assessed the cause of slow Ca^2+^ oscillation by adding a second round of CYN (20 min after a fast-to-slow mode switch). As shown in Figs. 3h and 3i, CYN reactivated the fast Ca^2+^ oscillation, and the islets recovered to slow oscillation after washout. Consistently, the application of CYN increased cAMP level (Fig. 3f) and stimulated endogenous glucagon secretion (Fig. 3g). These results indicated that the inhibition of α cells by δ cells led to insufficient glucagon secretion, resulting in slow Ca^2+^ oscillation in islet β cells.

Next, we studied how δ cells affect the α and β cell Ca^2+^ activities using *Glu-Cre^+^; GCaMP6f^f/+^; Ins2-RCaMP1.07* islets. α and β cells exhibited synchronized activation, with α cells activating 25 s after β cells. Adding CYN evoked fast α and β Ca^2+^ oscillation (Figs. 3j and 3k). However, the reduction in Ca^2+^ oscillation period did not affect the delay of α cells (22 s with CYN), which faithfully recapitulated the effect of adding glucagon^19^. Our findings suggested that CYN liberated α cells from δ cells, leading to fast islet Ca^2+^ oscillation.

### Optogenetic Inhibition and Relief of δ Cells Switch Islet between Fast and Slow Ca^2+^ Oscillations

To exert precise temporal control over δ cells, we employed *Sst-Cre^+/-^;Ai35^f/+^* mouse islets in which the δ cells expressed light-sensitive proton pumps (Fig. S3a)^29^. In these islets, with the 561 nm light being on or off, the δ cells can be turn off or on. During the 5-10 minutes light exposure, the islets switched from slow to fast oscillation with a period of approximately 20 s (Figs. 4a, 4c and Video 1). When the light was turned off, the islets returned to slow oscillation with a period of around 40-150 s. We fitted the Hill function to the time-dependent oscillation period change after turning on/off the light (Figs. 4b and 4d) and found that the period was reduced by an average of 69 s, with a delay of 126 s (light on). Conversely, when the light was turned off, the period increased by an average of 63 s, with a delay of 209 s. In control islets isolated from *Sst-Cre^+/-^ and Ai35^f/+^* mice, the 561 nm light had no significant effect or slightly prolonged the oscillation period (Figs. 4e and S3b). The optogenetic inhibition and relief of δ cells caused islets to switch between fast and slow oscillations. These results highlight the importance of δ cells to generate tunable islet oscillation patterns.

**Fig. 4.**
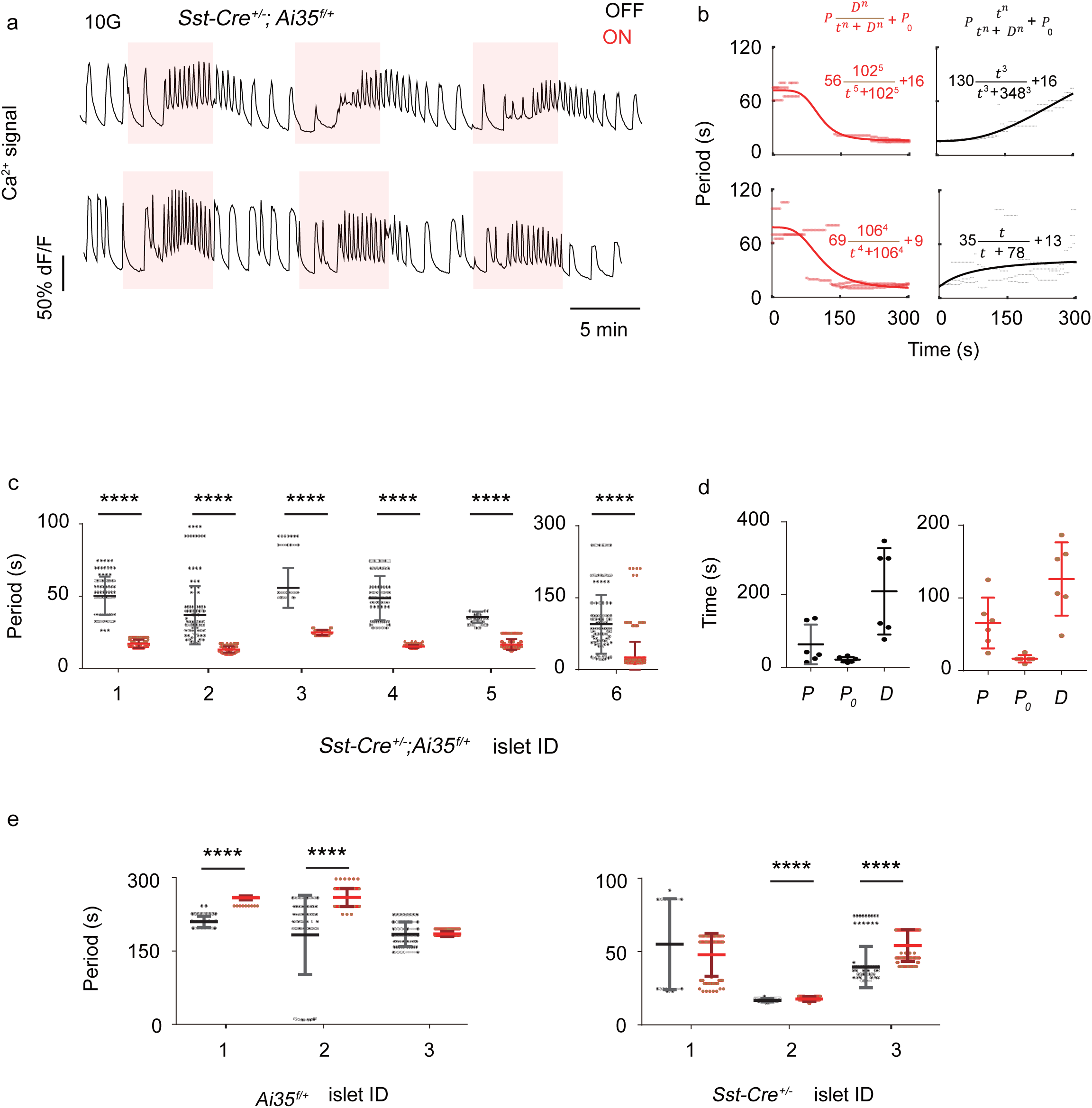
Optogenetic Inhibition of δ Cells Induces Fast Ca^2+^ Oscillation. a) Recordings of whole islet Ca^2+^ signals from *Sst-Cre*^+/-^;*Ai35*^f/+^ mouse islets. The red shadow shows the 5 min 561 nm light-on window (see Methods). b) Oscillation period with 561 nm light on (red dots) and off (black dots). The time turning light on/off is marked as 0 s. The red/black curves fit the light on/off periods with inhibitory/stimulatory hill function (see Methods). The top/bottom row corresponds to the top/bottom trace in (a). c) Oscillation period before (black) and after (red) 561 nm light on for 6 islets from *Sst-Cre*^+/-^;*Ai35*^f/+^ mice. Error bars in all panels depict SDs. d) Fitted parameters *P*, *P*_0_, and *D* in inhibitory (red)/stimulatory (black) hill function. e) Oscillation period before (black) and after (red) 561 nm light on for islets from *Ai35*^f/+^ (left) and *Sst-Cre*^+/-^ mice (right).

### Continuous Tuning of Parameters Reveals an Intermediate Oscillation Mode between Fast and Slow

To understand the conditions under which distinct Ca^2+^ oscillation modes occur, we applied gradient somatostatin concentrations (30, 70, and 100 nM) to islets with decreasing α cell number.

Under 10 mM glucose stimulation, the islets with abundant α cells showed stable fast oscillation, with a single peak distribution in the period (Figs. 5a and 5b, blue, mean value is ∼20 s). Elevated somatostatin concentration initially induced mixed oscillations of fast and slow, with a bimodal period distribution (Figs. 5a and 5b, yellow, mean values are ∼20 s and ∼180 s). Further increase in somatostatin concentration induced slow oscillation with single peak distribution (Figs. 5a and 5b, red, mean value is ∼200 s). This indicated that gradual increase in somatostatin can lead to two types of transition successively in the islet oscillation mode - fast to mixed and then to slow.

**Fig. 5.**
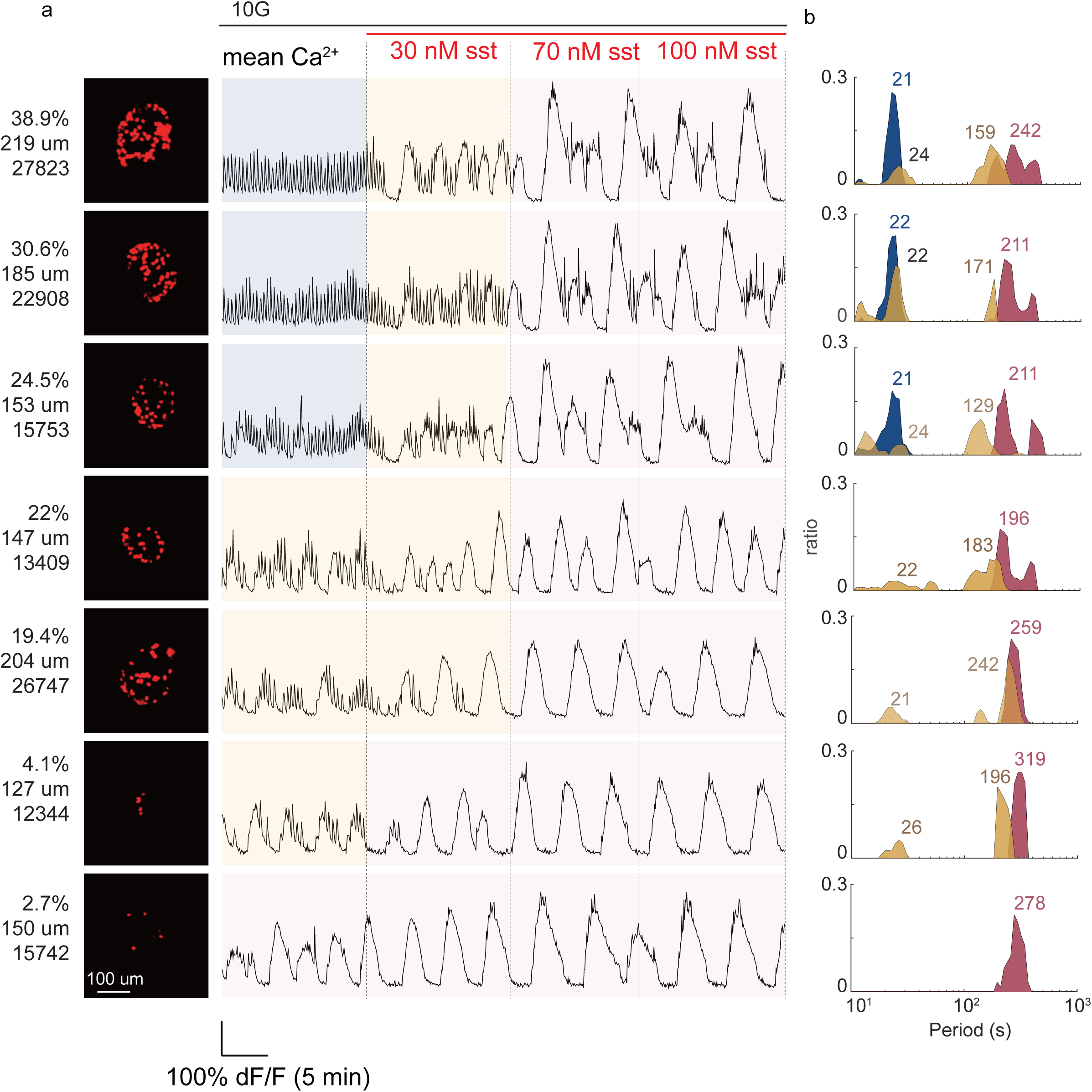
The Transition of Fast-Mixed-Slow Oscillations. a) Left panel: Maximum intensity projection of islets showing α cell (TdTomato) distribution. The numbers indicate the percentage of α cell area and the islet diameter. Islets are arranged from top to bottom based on decreasing α cell number. Right panel: Recordings of whole islet Ca^2+^ signal with 10G and different somatostatin concentration (30 nM, 70 nM and 100 nM somatostatin). Shadow represents the fast (blue), mixed (yellow) and slow (red) Ca^2+^ oscillation. b) Oscillation period distribution for islet showing fast (blue), mixed (yellow) and slow (red) Ca^2+^ oscillation. Each row corresponds to the same islet in a). Numbers indicate the peak Ca^2+^ oscillation period.

Interestingly, the same rule applies to the α cell number parameter. Under 10 mM glucose stimulation, islets with abundant α cells (> 24.5%) showed fast oscillation (a single peak distribution of period). However, islets with a moderate number of α cells (4% - 24.5%) exhibited mixed oscillations (a bimodal distribution of period). When the number of α cells is lower than 4%, islets showed pure slow oscillation (a single peak distribution of period). In summary, the islets exhibited fast, mixed and slow oscillations as the number of α cells in the islet deceased.

### A Mathematical Model of Islet Ca^2+^ Oscillation

The α cells secrete glucagon, stimulating hormone secretion in islet cells by elevating their cAMP level^17^. δ cells, on the other hand, release somatostatin, inhibiting islet cells’ hormone secretion by decreasing their cAMP level^23^. Through simultaneous cAMP/Ca^2+^ imaging, we observed a correlation between islet Ca^2+^ frequencies and the cAMP level controlled by glucagon and somatostatin (Figs.6a and 6b). Somatostatin inhibited cAMP level, leading to slow Ca^2+^ oscillation in the islet. In contract, forskolin and glucagon increased cAMP level, resulting in fast Ca^2+^ oscillation. Following washout, cAMP level recovered, and the islet returned to slow oscillation. Thus, we assumed that α and δ cells together control the islet Ca^2+^ oscillation modes by tuning cAMP levels (Fig. 6c). The model accounted for the opposing influence of α and δ cells on intracellular cAMP levels using a Hill equation 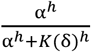. δ cells right shifted α cells’ activation threshold 𝐾(δ) (Fig. 2a), specifically using 𝐾(δ) = δ. Since we observed that islet’s Ca^2+^ oscillation periods exhibited a bimodal distribution (Figs. 2a and 5b), we assumed that the activity of β cells includes a fast oscillation part (𝜔_𝛼_) influenced by cAMP/glucagon and a slow oscillation part (𝜔) by default. The combination coefficient of these two parts is countered by somatostatin 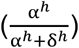. The activity of α and δ cells was governed by cAMP-dependent production, basal secretion, and degradation. The equations are as follows and Table 1:

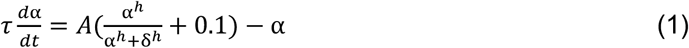

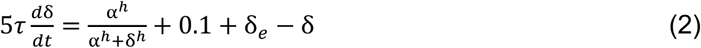

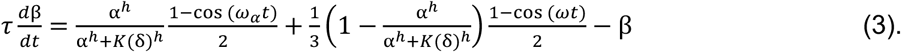

**Fig. 6.**
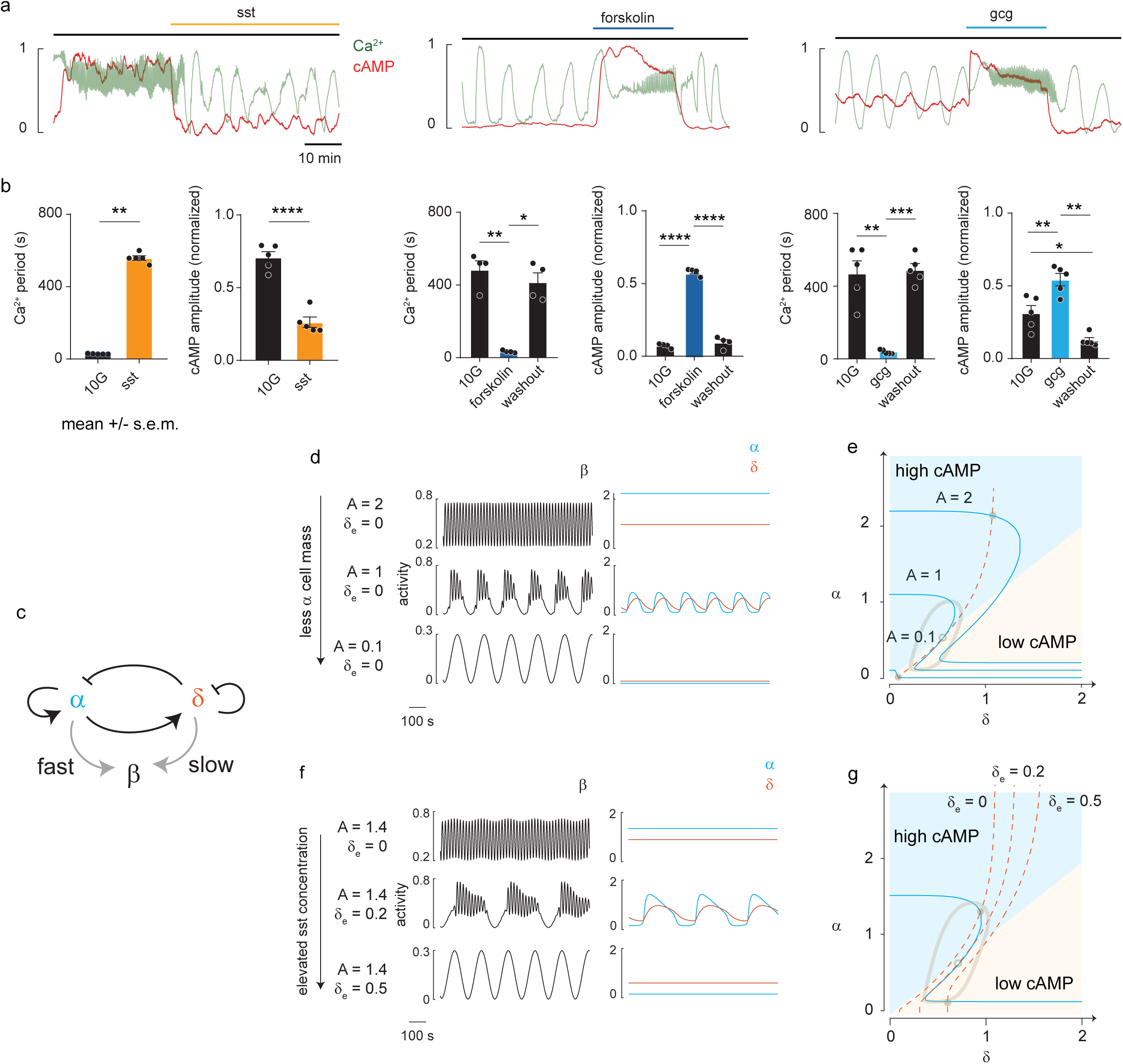
Mathematical Model of Islet Fast, Slow and Mixed Oscillation. a) Whole islet Ca^2+^ (green) and cAMP (red) signal before treatment, after 100 nM somatostatin, 120 nM forskolin and 100 nM glucagon perturbation. Islets were isolated from *Ella-Cre*^+^*;jRGECO1a^f/+^*mice. b) Mean Ca^2+^ oscillation period and mean cAMP amplitude before treatment, after somatostatin (n = 5 islets from 3 mice), 120 nM forskolin (n = 5 islets from 3 mice) and 100 nM glucagon (n = 4 islets from 3mice) perturbation. Error bars in all panels depict SEMs. c) Schematic of islet model. d) Activity traces of β cell (black, left panel), α cell (blue, right panel) and δ cell (yellow, right panel) with decreasing α cell mass. Parameter *A* for α cell mass and δ*_e_* for exogenous somatostatin. e) Phase planes of the islet model. Nullcline of α cell equation is denoted by blue line. Nullcline of the δ cell equation is denoted by dash yellow line. The intersections of the nullclines are the positions of the fixed points. When A is 2, the fixed point is stable (high cAMP state). When A is 1, the fixed point is unstable and surrounded by a stable limit cycle (oscillated cAMP state). When A is 0.1, the fixed point is stable (low cAMP state). f) Same as b with elevated somatostatin concentration (δ*_e_*). g) Same as c. When δ*_e_* is 0, the fixed point is stable (high cAMP state). When δ*_e_* is 0.2, the fixed point is unstable and surrounded by a stable limit cycle (oscillated cAMP state). When δ*_e_* is 0.5, the fixed point is stable (low cAMP state).

**Table 1.** Parameters value in mathematical model.

|  |  |
| --- | --- |
| $h$ | 5 |
| $\omega_{\alpha}$ | $\frac{2\pi}{20}$ |
| $\omega$ | $\frac{2\pi}{180}$ |
| $\tau$ | 10 |

The parameters 𝐴 and δ_𝑒_ represented α cell number and exogenous somatostatin concentration, respectively. Note that the system’s behavior mainly depends on the coupling between α and δ cells and their relative activity (i.e., cAMP level) characterizes β cell oscillation patterns.

By adjusting the α cell number (𝐴) or exogenous somatostatin concentration (δ_𝑒_), our model displayed a transition from fast to mixed to slow oscillation (Figs. 6d and 6f). This transition came from a Hopf bifurcation, which is the appearance and disappearance of a periodic orbit through a fixed point as parameter varies (Figs. 6e and 6g, fixed point - periodic orbit - fixed point). When the α cell was active (large 𝐴 or small δ_𝑒_), the model showed a high cAMP state fixed point with higher α cell activity than δ cell activity and the β cell exhibited fast oscillation (Figs. 6d and 6f, first row). When the α cell was inactive (small 𝐴 or large δ_𝑒_), the model presented a low cAMP state fixed point with lower α cell activity than δ cell activity (Figs. 6d and 6f, third row), and the β cell exhibited slow oscillation. When the α cell was moderately active, due to the coupling with the δ cell, the system showed an alternating high and low cAMP state, and the β cell exhibited mixed oscillations (Figs. 6d and 6f, second row). Here the model suggests that the mixed oscillation is a result of a balanced paracrine coupling between α and δ cells.

## Discussion

In a mouse islet, δ cells represent only a small percentage (∼5%), but they have an intriguing spatial distribution. δ cells are intermingled with α cells and occupy the periphery of the islet, enveloping the β cells in the core^30–32^. The present study demonstrates that δ cells play a role in tuning the oscillation modes. Our pharmacological and optogenetic experiments showed that the interaction between δ and α cells is the main origin of variability in islets’ oscillation patterns.

Combined with a model where β cells oscillate slowly by default, glucagon induces faster oscillations by elevating cAMP. This effect can be countered by somatostatin, which restores β cells’ default slow oscillations. This is consistent with a recent study showing that transit chemical inhibition of δ cell in intact islets accelerates oscillation^23^. We note that a recent paper showed islets with δ cell ablation oscillated slowly in response to glucose^33^. These slow Ca^2+^ oscillations appear to be the default state of β cells, independent of somatostatin (Fig.5 bottom islet).

Interestingly, despite the structural and compositional diversity of islets, the oscillation period presents a bimodal distribution with center peaks of ∼20 s and ∼200 s. Similar results have also been reported in previous studies^21,34,35^. Moreover, although there is apparent heterogeneity in Ca^2+^ activities across different islets, each individual islet exhibited a conserved Ca^2+^ oscillation pattern. The tight dependence on the α cell mass suggests that the diverse paracrine environment leads to distinct oscillation modes. The mechanisms by which continuous variations in structure elicit discrete activity patterns require further exploration.

Pharmacological perturbation of paracrine interactions causes bidirectional mode switching within the same islet. The addition of glucagon or blocking somatostatin (Sstr2 blocker CYN) leads to a switch from slow-to-fast mode. On the other hand, blocking glucagon (using Gcgr and Glp1r blockers, MK0893 and Ex9^19^) or adding somatostatin causes a switch from fast to slow mode. Induction of slow-to-fast switching is relatively easy (100 nM glucagon) and complete (independent of α cell mass) in the perturbation experiment. In contrast, inducing the fast-to-slow transition is relatively difficult (1 µM somatostatin), incomplete (dependent of α cell mass) and typically with a time delay. Moreover, by using the somatostatin sensor^36^, we observed that the endogenous somatostatin concentration can reach >1 uM (Fig. S1b). 100 nM somatostatin induced an incompetent transition from fast to slow/mixed oscillations and 1 uM somatostatin induced a larger slow spectrum (Fig. S1c and S1d). This suggests that islet cells could be tightly controlled by the accumulated somatostatin concentration – given that complete blocking is difficult and recovery experiments are easy.

Recent studies have found that somatostatin can directly affect the Ca^2+^ activity of β cells^23^, indicating that there are two levels of integration of δ-α signaling - the regulation of endogenous glucagon level in islets and the balance of cAMP level in β cells. Our study found that Sstr2 antagonists have a significant effect on islet Ca^2+^ activity. In contrast, the group treated with the Sstr3 antagonist MK4256 did not exhibit significant differences on oscillation frequency (see Figs. S2a and S2b). Interestingly, we observed that with 100 nM exogenous somatostatin, MK4256 increased the oscillation frequency (see Figs. S2c and S2d). This difference suggests an uneven endogenous somatostatin concentration. Moreover, the acceleration does not appear to be mediated via glucagon, as it persists even with the introduction of ex9 (see Figs. S2i and S2j). This suggests that the regulation of endogenous glucagon secretion is a primary integration mechanism. This is consistent with recent reports indicating that glucagon secretion is indirectly inhibited by somatostatin at high glucose levels, as revealed by K_ATP_ channel blocker^37^.

Our optogenetic interference revealed that a minority of δ cells can switch the majority of islet cells’ Ca^2+^ oscillation modes. An interesting question arises regarding the upstream signal of δ cells. Single-cell sequencing has found that δ cells have a unique receptor expression pattern^38,39^, including selective expression of the ghrelin receptor^26,38^. Further investigation using pharmacological perturbation and in vivo Ca^2+^ imaging studies are needed to better understand the role of δ cells and the design principle of the pancreatic islet^40,41^.

Previous studies have shown that glucagon promotes adenylyl cyclase-catalyzed generation, leading to elevated cAMP level^17,42^. Conversely, somatostatin reduces cAMP level through inhibiting adenylyl cyclase^43^. Stimulation of cAMP induces fast Ca^2+^ oscillation^44,45^. Thus, glucagon insufficiency, somatostatin accumulation, or a combination of both could cause the oscillation mode transition. In this study, we noticed the relieve of endogenous glucagon via CYN lead to a spontaneous fast-to-slow Ca oscillation modes switch (∼30 mins delay). Further studies could be conducted to comprehend the mechanism underlying that process.

Studies from 1994^8^ to 2022^23^ have consistently reported that large islets appeared to be more resistant to the loss of fast oscillation during pharmacological perturbation. In light of the heterogeneity observed in the somatostatin’s action in this study, it is likely that it is due to the large intra-islet α cell mass, rather than insufficient drug penetration.

## Acknowledgements

We are grateful to Erik Gylfe, Anders Tengholm and Palle Serup for helpful discussions. We thank Chunxiong Luo and Shujing Wang for their help with the microfluidic chip design. We thank the Imaging Core Facility and the State Key Laboratory of Membrane Biology for assistance with two-photon microscopy. We thank Dr. Ye Liang for providing the technical support of Imaris software. This work was supported by grants from the National Natural Science Foundation of China (12090053, 32088101), The National Key Research and Development Program of China (2018YFA0900700, 2021YFF1200500). H.R. was supported by the Boya Postdoctoral Fellowship of Peking University. B.X. was supported by the Postdoctoral Fellowship of Peking-Tsinghua Center for Life Science.

## Methods

### Animals

The study was conducted according to the guidelines of the Declaration of Helsinki and approved by the Ethics Committee of Peking University (protocol code #IMM-ChenLY-1). All animal experiments were conducted following the institutional guidelines for experimental animals at Peking University, which were approved by the AAALAC. Mice were housed with a 12-h on/12-h off light cycle, 20-24 ℃ ambient temperature and 40-70% humidity. *Ins1-Cre^+/-^; GCaMP6f^f/+^* mice were generated by crossbreeding *Ins1-Cre* mice (Jackson Laboratory, Strain #:026801) with *Rosa26-GCaMP6f^flox^* mice (Jackson Laboratory, Strain #:028865). Experiment shown in Fig.3a, 3b, 3d, 3e, 3h, 3i, S1a-c and S2a-j used islets isolated from *Ins1-Cre^+/-^; GCaMP6f^f/+^* mice. *Ella-Cre*^+^;*GCaMP6f^f/+^* mice were generated by crossbreeding *EIIa-Cre* mice (Jackson Laboratory, Strain #:003314) with *Rosa26-GCaMP6f^flox^* mice. *GCaMP6f^+^* mice were generated by multiply rounds crossbreeding of *EIIa-Cre^+^;GCaMP6f^f^* mice until spontaneous germline recombination was detected. The missing of stop codon between flox sites was confirmed by PCR using template tail DNA extracted from *EIIa-Cre^-^;GCaMP6f^f^* mice. *Glu-Cre^+^; tdTomato^f/f^* mice were generated by crossbreeding *Glu-Cre* mice^46^ with *Rosa26-tdTomato^flox^* mice (Jackson Laboratory, Strain #:007909). *Glu-Cre^+^; tdTomato^f/+^; GCaMP6f^+^* were generated by crossbreeding *Glu-Cre^+^; tdTomato^f/f^* mice with *GCaMP6f^+^* mice. Experiment shown in Fig.1b-c, 2a-d, 5a-b used islets isolated from *Glu-Cre^+^; tdTomato^f/+^; GCaMP6f^+^* mice. *Glu-Cre^+^; GCaMP6f^f/+^; Ins2-RCaMP1.07* mice has been reporter^19^. *Sst-Cre^+/-^;Ai35^f/+^* mice were generated by crossbreeding *Sst-Cre* mice (Jackson Laboratory, Strain #:013044) with *Rosa26-Ai35^flox^* mice (Jackson Laboratory, Strain #:012735). We confirmed the labeling accuracy by immunofluorescence (Fig. S4). Analysis of tdTomato-expressing cells revealed that 96.7% were positive for somatostatin (1st row, n = 123 cells), 8.1% were positive for insulin (2nd row, n = 37 cells) and 0% were positive for glucagon (3rd row, n = 118 cells). The labelling efficiency for somatostatin is 83% in *Sst-Cre^+/-^*; *tdTomato^f+f^* mice. *Ella-Cre*^+^*;jRGECO1a^f/+^* mice were generated by crossbreeding *EIIa-Cre* mice (Jackson Laboratory, Strain #:003314) with *Rosa26-jRGECO1a^flox^* mice (Cat. NO. NM-KI-215001, Shanghai Model Organisms Center, Inc..). All mice were genotyped by PCR using template tail DNA extracted by the TIANamp Genomic DNAKit (DP304, TIANGEN). Mice were PCR genotyped with 2x EasyTaq® PCR SuperMix (AS111, TRANSGEN). Genotyping primers and protocols are available at Jackson Laboratory’s website. Mice were maintained in one cage with a light/dark cycle of 12 h and administered chow diet ad libitum. Both female and male mice aged 8-20 weeks were used in the study.

### Immunofluorescence of single islet cells

Immunofluorescence experiment and analysis of *Sst-Cre* mice was performed as previously described^47,48^. *Sst-Cre^+/-^;tdTomato^f/+^* islets were dissociation into single-cells with 0.25% trypsin-EDTA (25300-062, Gibco) for 3 min at 37℃ followed by briefly shaking and stopped by the culture medium. The cells were centrifuged at 94 g for 5 min and suspended by culture medium. The cell suspension was plated on coverslips in the poly-L-lysine-coated Glass Bottom Dish (D35-14-1-N, Cellvis) and cultured overnight. Then the cells were fixed in 4% PFA (Paraformaldehyde Fix Solution) for 40 min before permeabilization in PBS, 0.3% Triton-100X for 2 h, and blocking in PBS, 5% BSA, 0.15% Triton-100X for 1 h at room temperature. The samples were incubated for 2 h at 4℃ in a guinea pig anti-insulin antibody (1:200, A0564, Dako), a rabbit anti-somatostatin antibody (1:500, GTX133119, GeneTex) and a mouse anti-glucagon antibody (1:500, G2654, Sigma) separately. The islets were then incubated for 1h at 4 ℃ with Goat anti-Guinea pig immunoglobulin G (IgG) (H+L) secondary antibody (DyLight^TM^ 488) (1:1000, SA5-10094, Invitrogen), Donkey Anti-Rabbit IgG H&L (Alexa Fluor 488) (1:1000, ab150073, abcam), Goat anti-Mouse IgG H&L (Alexa Fluor 488) (1:1000, A32766, Invitrogen).

### Islet Isolation

Mice were euthanized, and freshly-prepared collagenase P solution (0.5 mg/ml) was injected into the pancreas via the common bile duct. The perfused pancreas was digested at 37 °C for 15 min, and the islets were handpicked under a stereoscopic microscope. After isolation (defined as day 0), the islets were cultured overnight in RPMI-1640 medium (11879020, Gibco) containing 10% fetal bovine serum (10099141C, Gibco), 8 mM D-glucose, 100 unit/ml penicillin and 100 mg/ml streptomycin for overnight culture (generally at 5 p.m. at day 0). They were maintained at 37℃ and 5% CO_2_ in a culture incubator before the imaging experiments. Imaging experiments were performed on day 1 and 2 (10 a.m. – 10 p.m.). The time between islet isolation and the experiment typically were 17 – 53 h.

### Microfluidic Chips

Microfluidic chips were fabricated by using the elastomer polydimethylsiloxane (PDMS)^49^. Briefly, we used the photo-polymerizable epoxy resin (SU8 3050) to make a positive relief master, and the PDMS mold was cured on the master. PDMS mold was removed from the master as the channeled substrate. Then we used an air plasma treatment to bond the PDMS mold with a glass slide (70 x 105 x 10 mm, Xinxiang Vic Science & Education Co., Ltd.). The islet trapping region was designed as a stair-like channel, using five different thicknesses of SU-8 photoresist. The circular ports in the chip are color-coded according to their function (Fig. 1a). The chip features eight independent inlet ports for loading and trapping islets. The left input solutions are evenly separated into channels 1, 2, 3, and 4, while the right input solutions are evenly separated into channels 5, 6, 7, and 8. To trap islets of different sizes, the heights of this region were designed to be 40, 80, 110, 150 and 300 μm. Before imaging, we degassed the chip and all the solutions with a vacuum pump for 5 min to achieve stable hour-long imaging. The microfluidic chip was pre-filled with KRBB solution (125 mM NaCl, 5.9 mM KCl, 2.56 mM CaCl2, 1.2 mM MgCl2, 1 mM L-glutamine, 25 mM HEPES, 0.1% BSA, pH 7.4) containing 3 mM D-glucose before use. Then we injected the islet into the microfluidic chip using a 10 μL pipette from the islet inlet shown in Fig.1a. The degassed solutions were loaded into 10 ml syringes with needles 25G x 20 mm. The syringe connected to the chip through the PA single-lumen tube (0.45*0.8 mm, Shenzhen Kaist Medical Technology Co., LTD) with the reagent inlet. During the imaging, the islet in the microfluidic chip was kept at 37°C and 5% CO2 on the microscope stage during imaging. The reagents were automatically pumped into the microfluidic chip with a flow rate of 1600 µl/hour (each islet obtained a flow rate of 400 µl/hour) by the TS-1B syringe pump (LongerPump), which was controlled by software written in MATLAB. Such device enabled the simultaneous comparison of two sets of observations – each consisted of a flexible combination of up to five perfusion solutions.

### Fluorescent Imaging of Ca^2+^ Signal and 3D Structural Composition

We used a Dragonfly 200 series (Andor) with a Zyla4.2 sCMOS camera (Andor, 2048 x 2048 pixels, and each pixel size was 6.5 µm x 6.5 µm) and Fusion software. All channels were collected with a 10x/0.85 NA Microscope Objective (Warranty Leica HCX PL APO). Four fields of view were combined each time. Thus, the actual imaging field of view is 5.3 mm x 1.3 mm (the imaging zone of the chip is 5 mm x 1 mm). The sampling time resolution was 4.5 s. For 4% transmission of 488 nm illumination (200 mW), the exposure was set to 50 ms. A single bandpass filter with a center wavelength of 525 nm (band width 50 nm) was used to collect the GCaMP6f emission. We used Olympus FVMPE-RS Multiphoton Excitation Microscope (using dual pulsed two-photon far-red lasers for deep tissue imaging) with GaAsP PMTs (FV30-AGAPD). The signal was collected with a water immersion objective for multi-photon 25X/1.05, W.D. 2.0mm (XLPLN25XWMP2). The GCaMP6f signal was excited at laser’s wavelength 920 nm and collected through green barrier filter (495-540 nm). The tdTomato signal was excited at laser’s wavelength 1045 nm and collected through red barrier filter (575-645 nm). The *spots* function in Imaris9.7 was used for α cell identification and counting (with the parameter of 10 um cell diameter). In this study, we utilized a two-photon microscope to capture the 3D islet image in Fig.1b and quantify the number of α cells in Figs. 2a-d. For all the recording Ca^2+^ and cAMP signals, we used a spinning disk microscope, representing an optical section layer.

### Adenovirus Infection and cAMP Imaging

For cAMP imaging, the *Ella-Cre*^+^*;jRGECO1a^f/+^* mice islets were infected after isolation with recombinant adenoviruses (pAdeno-MCMV-G-Flamp2^50^) by 1 h exposure in 200 μl culture medium (approximately 4×10^6^ plaque forming units (PFU)/islet), followed by addition of regular medium and further culture for 16-20 h before use. All fluorescence images were acquired using spinning-disc confocal microscopy (10x objective, Dragonfly) at time resolution of 4.5 s per frame. Note the cAMP sensor driven by CMV promoter was introduced to islets by adenovirus, without specificity for cAMP sensor expression. The cAMP signal represented the mean fluorescent intensity changes from the sensor-positive cells across the whole islet. Considering the cell ratio, we interpret the cAMP signal as originating from β cells.

### Hormone Concentration Detection

The effluent from each channel of microfluidic ship was continuously collected every 15min and frozen at -20℃. Insulin was measured using a mouse insulin AlphaLISA Kit (product No.:AL3184, PerkinElmer) or an insulin ELISA kit (10-113-01, Mercodia AB, Sweden). Glucagon was measured using a glucagon ELISA kit (10-1281-01, Mercodia AB, Sweden).

### Oscillation Spectrum Analysis

#### CWT analysis

CWT (continuous wavelet transformation) analyzes signals jointly in time and frequency. The continuous wavelet transformation compared a signal with shifted and scaled copies of a basic wavelet. We used the analytic Morse (3,60) wavelet with period limits from 10 s to 600 s as the continuous wavelet transform filter bank. The number of voices per octave (a doubling) was 10 (interpolate 10 frequencies every period doubling). The continuous wavelet transformation returned a matrix *cwt* of 60 rows and *T* columns of an input time series with *T* timepoints. Each row of *cwt* represented the spectrum for a specific frequency, such as row 1 for 10 s, row 11 for 20 s, row 21 for 40 s, row 60 for 600 s and row 27 for 60 s.

#### Removing edge-effect artifacts

The cone of influence region in matrix *cwt* affected by edge-effect artifacts was set zero (effects arise from areas where the stretched wavelets extend beyond the edges of the observation interval). The processed matrix *cwt* was an accurate time-frequency representation of the data (second row of Fig. S1a).

#### Spectral distribution of the input time series

Matrix *cwt* was further binarized and used for subsequent analysis (third row of Fig. S1a). The sum of each row of *cwt* gave the spectral distribution of the input time series. As it’s shown in the right panel of the second row of Fig. S1a. The spectral distribution typically followed a bimodal distribution – with a fast peak near 18 s and a slow peak near 242 s.

## Notes

### Competing Interest Statement

The authors have declared no competing interest.

### Summary of Updates

Author order has been corrected. Previously we wrongly place the corresponding author as the first place which should be the last one.

